# Survey of Climate-structured Mycobiomes in Staple Maize: Implications for Endemic Keshan and Kashin–Beck Diseases

**DOI:** 10.64898/2026.04.03.716289

**Authors:** Yingxue Wang, Kunyu Zhang, Yu Sun, Likun Yang, Jinwu Yang, Xiaocheng Wang, Yihong Wan, Guoping Xi, Liuliu Guo, Shuqiu Sun

## Abstract

Keshan disease (KD) and Kashin–Beck disease (KBD) are geographically restricted disorders in rural China with overlapping environmental and dietary risk factors. Selenium deficiency alone cannot explain their regional heterogeneity. Maize, a dietary staple in endemic areas, represents a key exposure pathway for climate-sensitive foodborne fungi and their metabolites. We profiled maize-associated fungal communities from seven villages across KD-, KBD-, KD-KBD co-endemic, and non-endemic regions using ITS sequencing and integrative bioinformatics. Fungal diversity, composition, trophic structure, and predicted biosynthetic gene cluster potential differed markedly among regions. KD-endemic areas were enriched in saprotrophic taxa such as *Penicillium* and *Aspergillus*, KBD-endemic regions favored cold- and humidity-adapted fungi, and KD-KBD co-endemic areas exhibited the highest predicted mycotoxin potential. Fungal patterns were strongly associated with regional temperature and humidity. These findings support a climate-sensitive, foodborne exposome framework, suggesting that variation in maize-associated fungi may contribute to endemic disease risk and highlighting the need for fungal surveillance in public health strategies.

**Graphical abstract:** 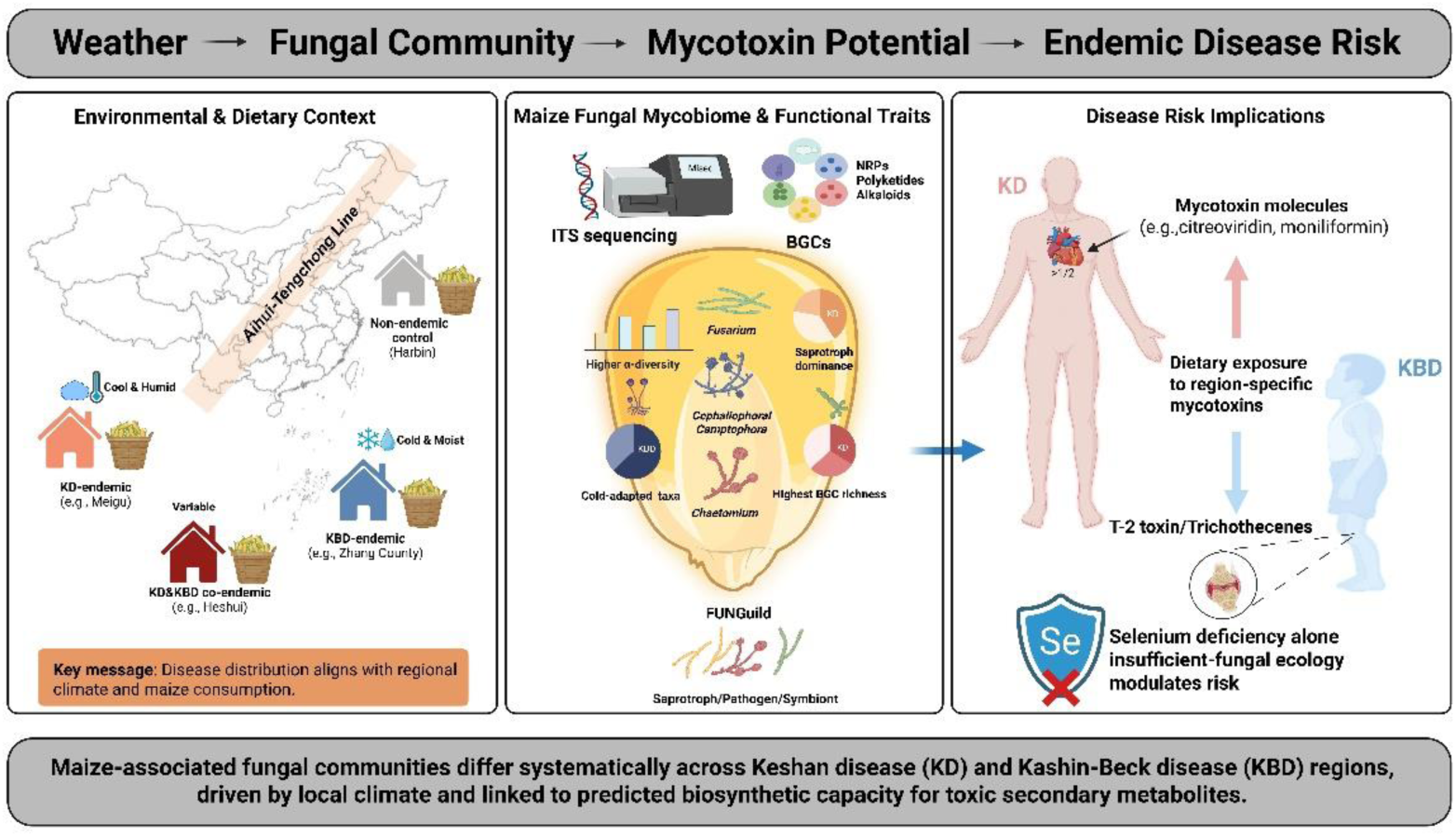

## 1. Introduction

Keshan disease (KD) and Kashin–Beck disease (KBD) are long-recognized endemic disorders affecting rural populations in China(Allander 1994; Li et al. 1985; Sun 2018). Despite decades of investigation, their etiologies remain incompletely resolved (Fang et al. 2012; Wang et al. 2013). While their clinical manifestations differ substantially—KD is characterized by multifocal myocardial necrosis and heart failure (Lei et al. 2011; Li et al. 1985), whereas KBD is a chronic osteochondropathy involving articular cartilage and growth plates—the two share striking epidemiological similarities (Lin et al. 2024; Wang et al. 2020; Zou et al. 2009). Both exhibit strong geographic clustering, a reliance on locally produced staple foods, and an association with impoverished rural environments, leading to their designation as “sister diseases” in mainland China (Wang et al. 2013).

In China, these diseases are distributed primarily along the Aihui-Tengchong Line, a region characterized by low soil selenium levels (Wang et al. 2013). Historically, etiological research focused on selenium deficiency, supported by evidence linking low selenium concentrations in soil and cereals to increased disease incidence (Prabhu and Lei 2016). Large-scale supplementation programs achieved partial success, particularly in reducing acute KD cases (Chen 2012). However, critical inconsistencies persist. “Healthy islands” remain within severely selenium-deficient areas, and selenium deficiency alone cannot explain why KD and KBD overlap in northeastern China but segregate in similarly deficient regions of the southwest (Yang 2013). These observations suggest that additional environmental or biological co-factors are required to trigger pathogenesis.

Mycotoxins—toxic secondary metabolites produced by filamentous fungi—have emerged as plausible contributors to the pathogenesis of both diseases (Guo 1986; Sun 2010; Yang 2002). Experimental models demonstrate that specific mycotoxins can induce pathological changes mimicking these disorders: moniliformin and citreoviridin cause myocardial injury analogous to KD, while T-2 toxin and related trichothecenes induce the chondrocyte necrosis characteristic of KBD (Chasseur et al. 1997; Li et al. 2026; Lu et al. 2021; Sun et al. 2019; Yu et al. 2022). These findings are reinforced by epidemiological data showing elevated mycotoxin contamination in staple grains, particularly maize, from endemic regions (Zhang 2012). Such associations suggest that chronic dietary exposure, resulting from the consumption of inadequately stored, fungus-contaminated cereals, may be a primary driver of disease prevalence.

Despite the evidence implicating mycotoxins, significant knowledge gaps remain regarding the ecological drivers of KD and KBD. Previous studies have largely focused on individual mycotoxins or isolated fungal strains, overlooking the complexity of the total mycobiome colonizing staple cereals (Sun 2012). Fungal community structure and the subsequent biosynthesis of toxins are shaped by a complex interplay of climatic conditions (temperature and humidity), agricultural practices, and interspecific interactions (Yan et al. 2026). Furthermore, the distinct clinical phenotypes of KD and KBD suggest the involvement of disease-specific fungal assemblages or unique toxigenic profiles (Yang 2013).

## 2. Experimental Procedures

### 2.1 Study Sites and Maize Sample Collection

All study sites were selected in accordance with official Chinese national standards and public health guidelines for the classification and management of endemic diseases, including GB 17020–2010, GB 16395–2011, and the *Three-Year Action Plan for the Prevention and Control of Endemic Diseases (2018–2020)*. Based on historical disease occurrence, study regions were classified into four categories: KD-endemic areas, KBD-endemic areas, co-endemic areas for KD and KBD, and non-endemic control areas.

KD-endemic sites included Meigu County (Sichuan Province) and Jingchuan County (Gansu Province). KBD-endemic sites were selected from Zhang County and Kangle County (Gansu Province). Co-endemic sites for KD and KBD were located in Heshui County and Zhengning County (Gansu Province). A non-endemic control site was selected in Pingfang District, Harbin City, Heilongjiang Province, which lies within the broader KD-KBD belt but has no documented history of either disease.

Maize kernels were collected during 2019–2020 from household-stored grain harvested in the preceding year. Samples were obtained from two KD-endemic villages (Meigu and Jingchuan), two KBD-endemic villages (Zhang and Kangle), two KD-KBD co-endemic villages (Heshui and Zhengning), and one non-endemic control village (Pingfang District, Harbin). In each village, five households were sampled, yielding a total of 35 maize samples. Approximately 500 g of maize kernels were collected from each household, placed into sterile kraft paper bags, labeled with unique identifiers, and transported to the laboratory under dry conditions for subsequent processing.

Detailed meteorological data for each study site during the sampling season, including mean temperature and maximum relative humidity, are provided in Supplementary Table 1. Sample collection was conducted during the local harvest season to ensure representative exposure of household-stored maize to environmental fungal colonization.

### 2.2 Laboratory Incubation of Maize for Fungal Enrichment

To promote fungal growth under controlled conditions, 60 g of maize kernels from each household sample were transferred into sterile, autoclaved Erlenmeyer flasks and incubated in a programmable environmental chamber (MJL-150; Tianjin Labotery Instrument Equipment Co., Ltd., Tianjin, China). Incubation conditions were designed to simulate typical local grain storage environments in endemic regions, with alternating diurnal temperatures of 25 °C during the day and 15 °C at night, and relative humidity maintained at 85 %. Incubations were conducted for approximately two weeks. Temperature and humidity were monitored daily to ensure stability. Incubation was terminated when visible fungal mycelial growth was consistently observed for 2–3 consecutive days, but before complete kernel colonization occurred, thereby minimizing overgrowth by fast-growing taxa.

### 2.3 Fungal DNA Extraction

Fungal mycelia were aseptically collected from the surface of incubated maize kernels, and approximately 0.15 g of material from each sample was immediately frozen and pulverized in liquid nitrogen using sterile, single-use mortars and pestles. Genomic DNA was extracted using a cetyltrimethylammonium bromide (CTAB)–based protocol. Briefly, homogenized samples were incubated in CTAB extraction buffer at 65 °C for 1 h, followed by three successive extractions with chloroform:isoamyl alcohol (25:1, v/v). DNA was precipitated with ice-cold isopropanol at -20 °C overnight, washed with 75 % ethanol, air-dried, and resuspended in TE buffer containing RNase A. DNA concentration and purity were assessed using a NanoDrop 2000 spectrophotometer (Thermo Scientific, USA), and all extracts were stored at -20 °C until downstream analyses.

### 2.4 PCR Amplification and ITS Sequencing

PCR amplification and ITS sequencing were performed by Allwegene BioTech Co., Ltd. (Beijing, China). The fungal ITS1 region was amplified using the primer pair ITS1-F (5′-CTTGGTCATTTAGAGGAAGTAA-3′) and ITS2 (5′-TGCGTTCTTCATCGATGC-3′). Each 25 μL PCR reaction contained 30 ng of template DNA, 5 μM of each primer, bovine serum albumin (BSA), and TransStart FastPfu DNA Polymerase Master Mix. Thermal cycling conditions consisted of an initial denaturation at 95 °C for 5 min, followed by 32 cycles of denaturation at 95 °C for 45 s, annealing at 55 °C for 50 s, and extension at 72 °C for 45 s, with a final extension at 72 °C for 10 min.

PCR products were verified by agarose gel electrophoresis, quantified, and pooled in equimolar concentrations. Sequencing libraries were constructed through Y-adapter ligation, bead-based purification, and PCR enrichment, followed by paired-end sequencing (2 × 250 bp) on an Illumina MiSeq platform using standard sequencing-by-synthesis chemistry.

### 2.5 Processing and Quality Control of Sequencing Data

Raw paired-end sequencing reads were subjected to stringent quality control to ensure high-confidence downstream analyses. Low-quality bases with Phred scores below 20 were removed, and a sliding-window trimming approach (50 bp window) was applied to discard read regions with average quality scores below 20. Reads shorter than 50 bp after trimming were excluded. High-quality paired-end reads were subsequently merged based on overlapping regions to generate raw tags. Chimeric sequences and abnormally short fragments were identified and removed, yielding a final set of clean tags for subsequent bioinformatic analyses.

### 2.6 OTU Clustering and Taxonomic Annotation

High-quality clean tags were clustered into operational taxonomic units (OTUs) at a 97% sequence similarity threshold using the UPARSE pipeline. Representative sequences from each OTU were taxonomically assigned using the UNITE ITS reference database in combination with the Ribosomal Database Project (RDP) classifier. To minimize the influence of sequencing artifacts and spurious taxa, OTUs with extremely low abundance or those present in fewer than two samples were removed. The resulting curated OTU table was used for all subsequent analyses of fungal diversity, community composition, and functional inference.

### 2.7 Alpha and Beta Diversity Analyses

Alpha diversity metrics, including the Shannon diversity index and Chao1 richness estimator, were calculated using the *phyloseq* package in R. Differences in alpha diversity among groups were assessed using pairwise Wilcoxon rank-sum tests, with p-values adjusted for multiple comparisons using the Benjamini–Hochberg false discovery rate (FDR) method. Beta diversity was evaluated based on Bray–Curtis dissimilarity matrices and visualized using principal coordinate analysis (PCoA). Statistical differences in community composition among groups were tested using permutational multivariate analysis of variance (PERMANOVA; *adonis2* function, 999 permutations).

### 2.8 Fungal Trophic Mode Annotation

Fungal ecological guilds and trophic modes, including saprotrophic, pathotrophic, symbiotrophic, and mixed strategies, were inferred using the FUNGuild database. Only annotations classified as “probable” or “highly probable” were retained to ensure reliability of functional assignments. Trophic mode distributions were subsequently summarized and compared across regions to assess functional differences in fungal community structure.

### 2.9 Differential Abundance Analysis

Region-specific differences in fungal community composition were examined at the genus level using complementary multivariate and differential abundance approaches. Hierarchical clustering and heatmap visualization based on Bray–Curtis dissimilarities were applied to illustrate overall compositional patterns across regions. Differentially abundant taxa were identified using linear discriminant analysis effect size (LEfSe), with an LDA score threshold of 2.0 applied to define discriminative features. Results from clustering and LEfSe analyses were integrated to identify robust region-specific fungal signatures.

### 2.10 Prediction of Biosynthetic Gene Cluster (BGC) Potential

The biosynthetic potential of fungal communities was inferred by mapping OTU-associated genera to a curated reference database compiled from antiSMASH and the Minimum Information about a Biosynthetic Gene cluster (MIBiG) repository. Predicted biosynthetic gene cluster (BGC) classes included polyketides, nonribosomal peptides (NRPs), terpenes, alkaloids, saccharides, and mixed clusters. Comparative analyses of BGC distributions across regions were visualized using bubble plots generated in R.

### 2.11 Statistical Analysis

Statistical analyses were performed using R software. Differences in alpha diversity indices, trophic mode proportions, and predicted BGC abundances among regions were evaluated using Wilcoxon rank-sum tests, with p-values adjusted for multiple comparisons using the Benjamini–Hochberg false discovery rate method. Differences in beta diversity were assessed using PERMANOVA based on Bray–Curtis dissimilarities. Differentially abundant genera were identified using LEfSe, as described above. Heatmaps, ordination plots, and bubble plots were generated using standard R visualization packages. Statistical significance was defined as *p* < 0.05.

## 3. Results

### 3.1 Comparison of Fungal Richness and Diversity Across Different Regions

To evaluate differences in fungal community richness, diversity, and structure among the four regions, sequencing depth and taxonomic coverage were first assessed using rarefaction analyses. Operational taxonomic unit (OTU) rarefaction curves for all samples approached saturation, indicating sufficient sequencing depth and comparable sampling effort across groups (Fig. 1A). Similarly, Shannon diversity rarefaction curves reached clear plateaus, suggesting that the majority of fungal diversity present in the samples was effectively captured (Fig. 1B). Venn diagram analysis revealed both shared and region-specific taxa among the four groups: a total of 578 OTUs were common to all regions, while each group also harbored unique OTUs, reflecting distinct regional fungal assemblages (Fig. 1C).

**FIGURE 1.**
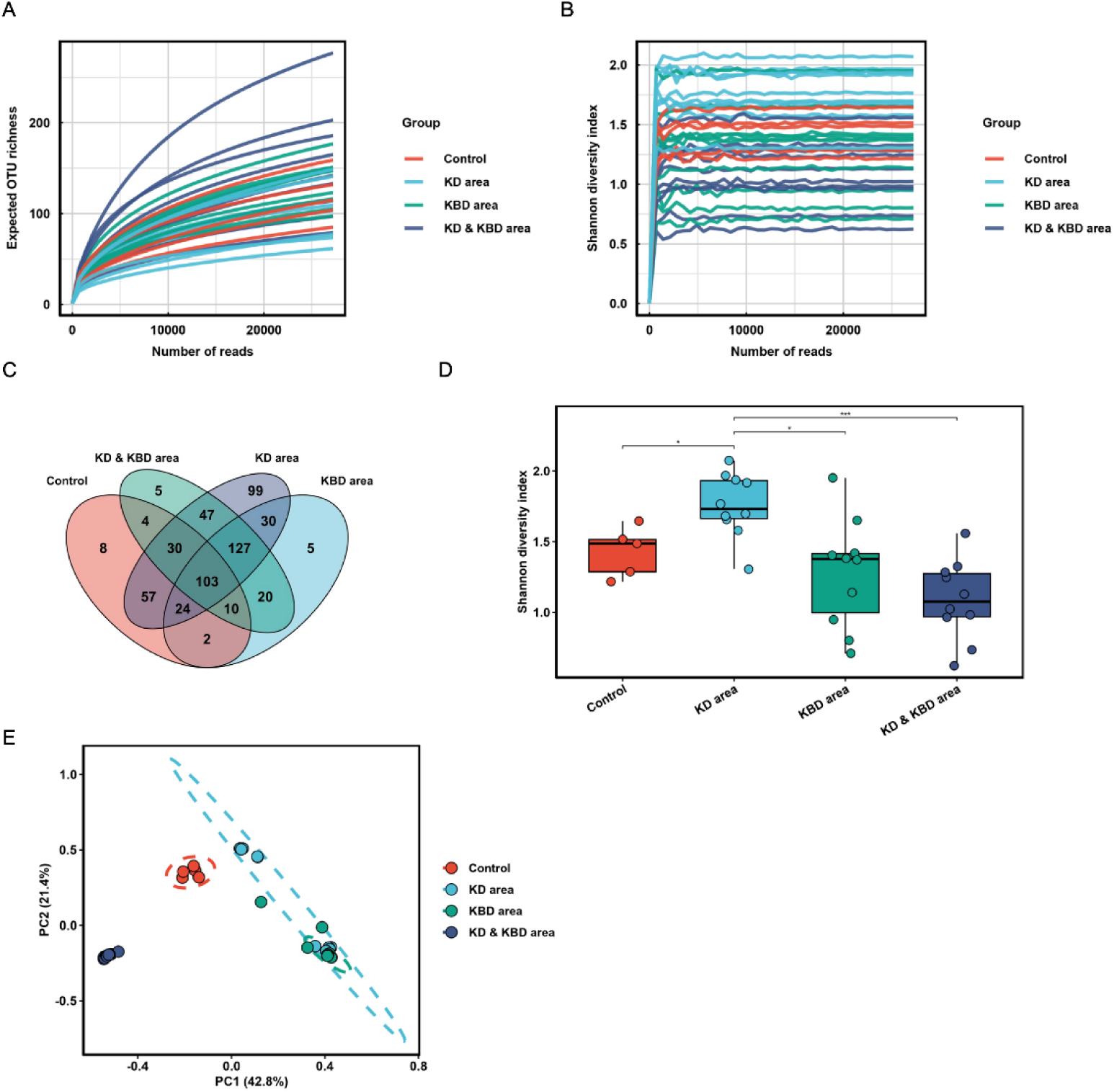
Comparison of fungal richness and diversity across different regions. (A) OTU rarefaction curves showing the relationship between sequencing depth and observed richness across the four groups. (B) Shannon diversity rarefaction curves indicating stabilization of fungal diversity with increasing sequencing depth. (C) Venn diagram illustrating shared and unique OTUs among the four groups. (D) Boxplots of the Shannon index comparing fungal diversity across regions. (E) Principal coordinate analysis (PCoA) based on Bray–Curtis distances showing distinct clustering of fungal communities among the four regions. \**p* < 0.05, \*\**p* < 0.01, \*\*\**p* < 0.001.

Prior to grouping by disease area, an initial assessment of the seven individual sampling sites revealed significant heterogeneity in both Shannon diversity and community structure (Supplementary Fig. 1A–B). Subsequent alpha diversity analyses of the four aggregated regions demonstrated significant differences in fungal richness and diversity. The Shannon index indicated that fungal richness varied significantly among groups, with the KD area exhibiting markedly higher richness than both the Control and KD & KBD areas (Fig. 1D), suggesting that KD-associated environments support more complex fungal communities. Beta diversity analysis based on Bray-Curtis dissimilarities further revealed clear separation of fungal community structures among the four groups (Fig. 1E). Samples from the Control formed a distinct cluster, clearly segregated from those of the KD, KBD, and KD & KBD areas, indicating pronounced compositional divergence. Collectively, these results demonstrate substantial geographic variation in fungal community richness, diversity, and overall structure across the four regions.

### 3.2 Fungal Community Composition and Trophic Functional Differences Across Regions

The taxonomic composition of fungal communities differed markedly across the four regions. At the phylum level, fungal assemblages were dominated by Ascomycota and Basidiomycota, a pattern consistently observed across the seven individual sampling sites (Supplementary Fig. 2A), although their relative abundances varied substantially among the aggregated groups (Fig. 2A). Notably, samples from the KD, KBD, and KD & KBD areas exhibited higher proportions of Basidiomycota compared with those from the Control region. At the genus level, pronounced geographic differentiation in community composition was evident (Fig. 2B), reflecting the distinct taxonomic profiles of the underlying sampling sites (Supplementary Fig. 2B). Several dominant taxa displayed region-specific enrichment patterns. Specifically, Fusarium, Stilbocrea, and Meyerozyma predominated in the Control group; Cephaliophora, Camptophora, and Russula were enriched in the KD area; Russula, Chaetomium, and Candida characterized the KBD area; and Cephaliophora, Camptophora, and Chaetomium were prominent in the KD & KBD area. These distinct taxonomic profiles underscore strong regional structuring of maize-associated fungal communities.

**FIGURE 2.**
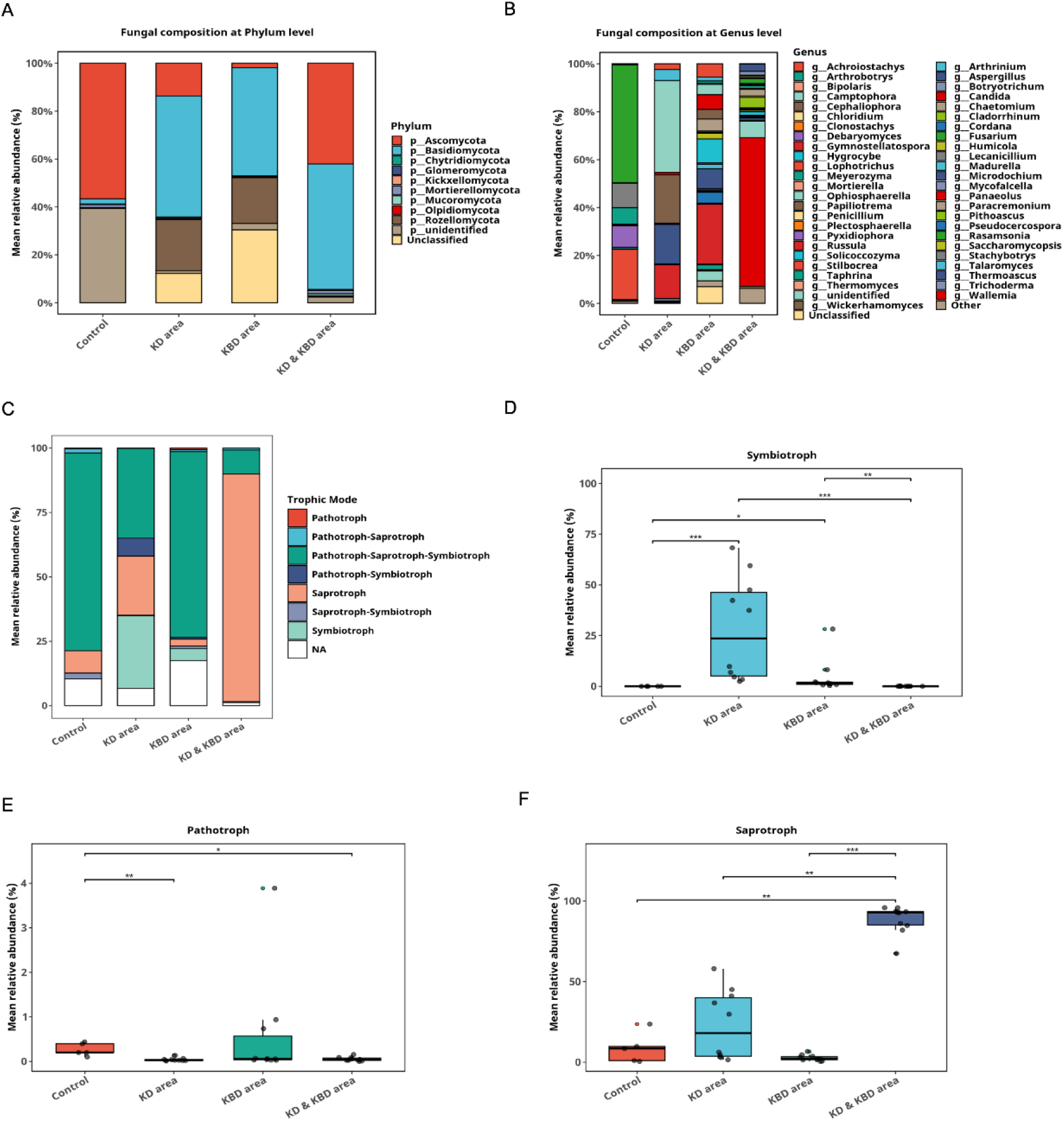
Fungal community composition and trophic functional differences across regions. (A) Relative abundance of major fungal phyla across the four groups. (B) Relative abundance of dominant fungal genera across the four groups. (C) Distribution of fungal trophic modes (symbiotrophs, pathotrophs and saprotrophs) among regions. (D–F) Boxplots comparing the relative abundances of symbiotrophs (D), pathotrophs (E), and saprotrophs (F) across regions. \**p* < 0.05, \*\**p* < 0.01, \*\*\**p* < 0.001.

Functional annotation based on fungal trophic modes further revealed substantial ecological differentiation among regions. The relative contributions of major trophic categories-including saprotrophs, pathotrophs, and symbiotrophs-varied significantly across groups (Fig. 2C). Quantitative comparisons showed that symbiotrophs were significantly enriched in the KD area relative to the other regions (Fig. 2D). Pathotroph abundance also differed markedly among groups, with the Control area exhibiting significantly higher levels (Fig. 2E). In contrast, saprotrophs reached their highest relative abundance in the KD & KBD area (Fig. 2F). Additionally, guilds exhibiting mixed trophic strategies (e.g., pathotroph-symbiotrophs) also displayed significant variations in abundance across the four groups (Supplementary Fig. 3). Collectively, these functional profiles suggest distinct fungal–host interaction potentials in different regions.

### 3.3 Differential Abundance and Region-Specific Fungal Signatures

To identify region-specific fungal signatures, differential abundance analysis was performed at the genus level. Heatmap visualization revealed pronounced heterogeneity in genus-level composition across the four regions (Fig. 3A). Hierarchical clustering based on Bray–Curtis distances showed that samples grouped primarily according to geographic origin, indicating strong spatial structuring of fungal communities. Several genera exhibited clear region-specific enrichment, with multiple taxa displaying significantly higher relative abundances in the KD, KBD, or KD & KBD areas compared with the Control group.

**FIGURE 3.**
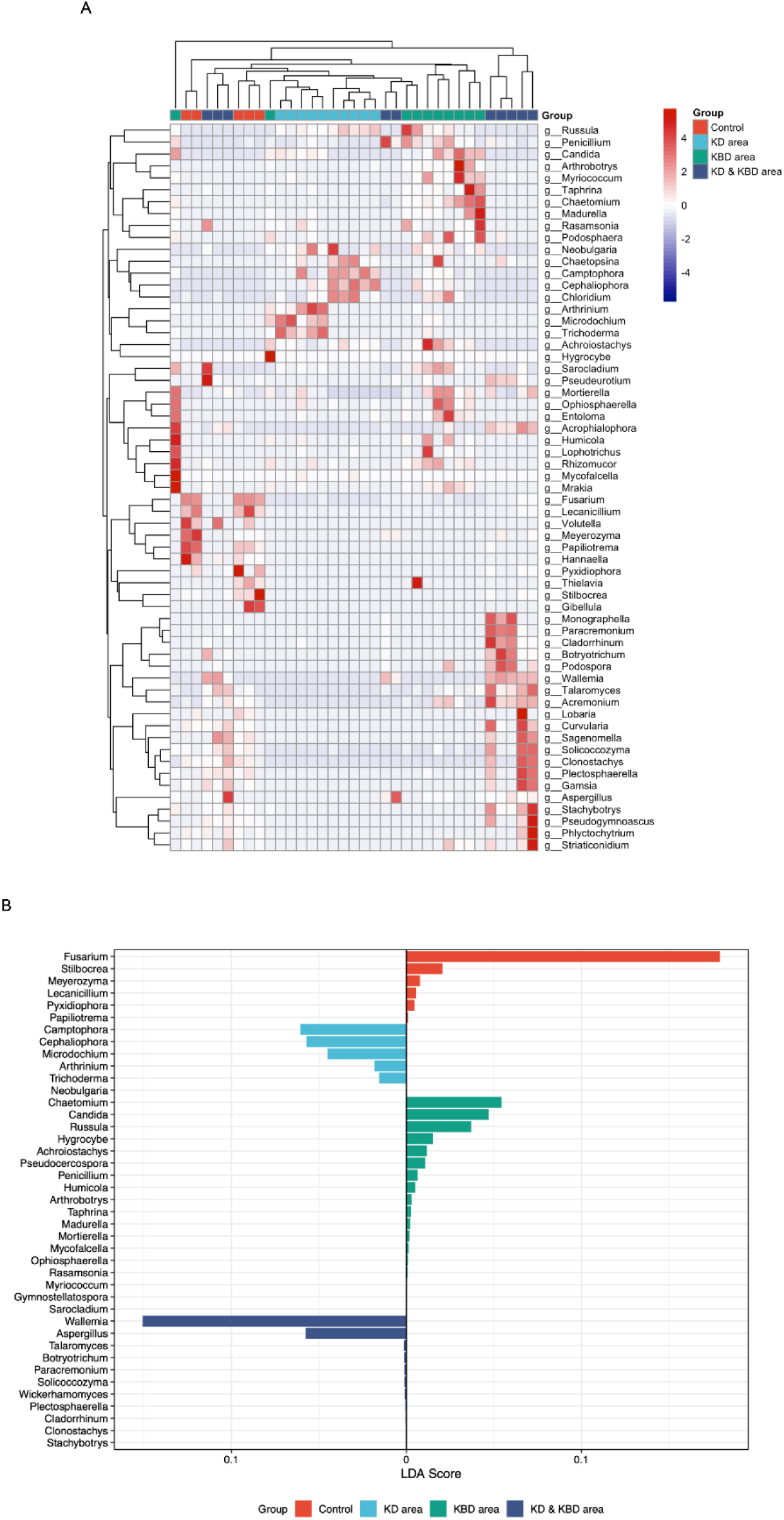
Differential abundance and region-specific fungal signatures. (A) Heatmap of the relative abundances of dominant fungal genera across the four groups. Hierarchical clustering based on Bray–Curtis distances reveals distinct region-specific distribution patterns. (B) Linear discriminant analysis effect size (LEfSe) identifying fungal genera that significantly discriminate among the four groups. Bars represent taxa with LDA scores above the predefined threshold and are colored according to the group in which they are enriched.

Linear discriminant analysis effect size (LEfSe) further identified key discriminatory genera contributing to compositional divergence among regions (Fig. 3B). The Control group was characterized by significant enrichment of *Fusarium*, *Stilbocrea*, and *Aspergillus*, reflecting the baseline fungal assemblage. In contrast, the KBD area was distinguished by *Russula*, *Chaetomium*, and *Candida*, whereas the KD area exhibited unique enrichment of *Cephaliophora* and *Camptophora*. The KD & KBD area also displayed distinct fungal signatures, with genera such as *Chaetomium*, *Microdochium*, and *Camptochaeta* showing strong discriminatory power. Collectively, these findings highlight pronounced regional differentiation in fungal community composition and identify taxa that may serve as potential biomarkers of region-specific ecological conditions.

### 3.4 Predicted Biosynthetic Potential of Fungal Communities Across Regions

Analysis of predicted biosynthetic gene clusters (BGCs) revealed pronounced regional differences in the metabolic potential of maize-associated fungal communities (Fig. 4). The distribution of BGC classes varied substantially among dominant genera, encompassing pathways associated with alkaloid, nonribosomal peptide (NRP), polyketide, terpene, saccharide, and mixed biosynthetic processes. Among the four regions, the KD & KBD area exhibited the highest overall predicted biosynthetic capacity, as indicated by larger bubble sizes and a broader diversity of BGC classes, particularly within genera such as *Aspergillus*, *Penicillium*, and *Pseudomonas*.

**FIGURE 4.**
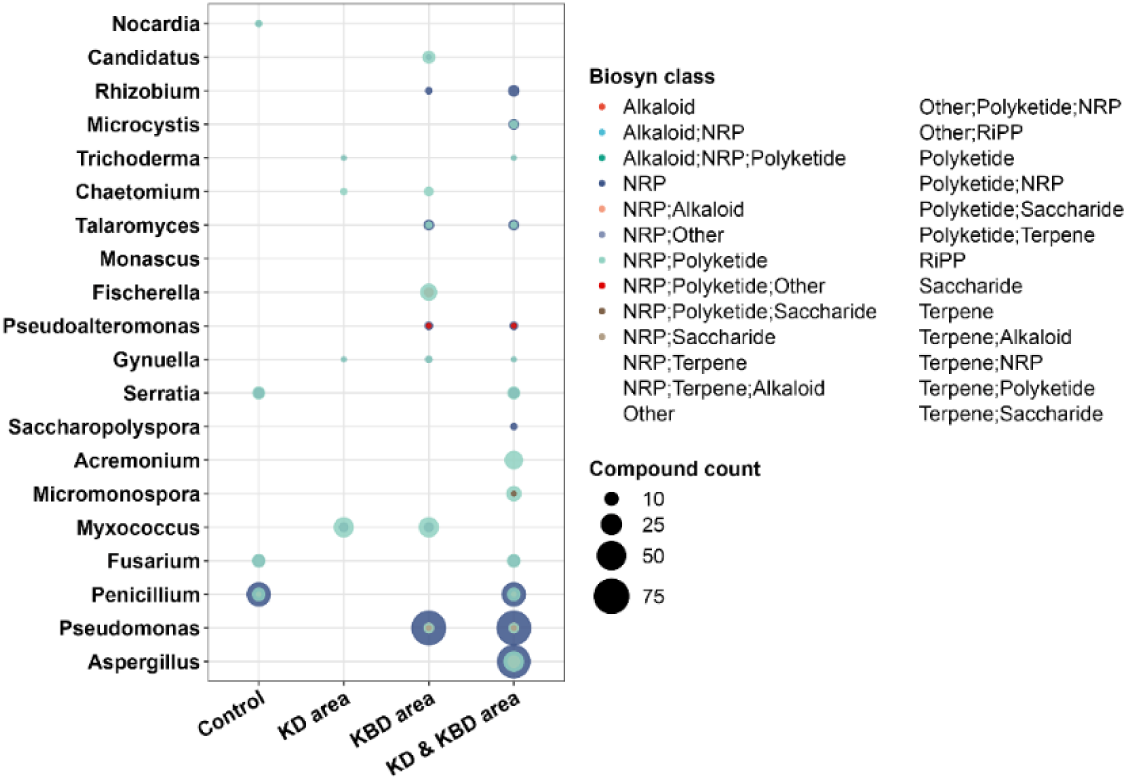
Predicted biosynthetic potential of fungal communities across regions. Bubble plot illustrating the distribution of predicted biosynthetic gene cluster (BGC) classes among dominant fungal genera across the four groups.

In contrast, fungal communities in the KD area displayed a more restricted yet distinct biosynthetic repertoire, largely associated with genera including *Micromonospora*, *Chaetomium*, and *Talaromyces*. The Control group showed the lowest diversity and abundance of predicted BGCs, suggesting limited secondary metabolic potential, whereas the KBD area exhibited intermediate biosynthetic capacity, driven primarily by genera such as *Fusarium*, *Trichoderma*, and *Saccharopolyspora*. Collectively, these results indicate that fungal communities in KD-associated regions possess greater functional versatility and an enhanced potential for secondary metabolite production compared with those in non-KD regions.

## 4. Discussion

Understanding the environmental determinants underlying the complex and historically dynamic geographic distributions of KD and KBD has remained a longstanding challenge (Yang 2000). While selenium deficiency is widely recognized as a major etiological factor, it fails to account for several persistent epidemiological inconsistencies, including the coexistence of KD and KBD in northeastern China, their segregation in selenium-deficient regions of southwestern China, and the presence of severely selenium-depleted areas without reported disease (Yang 1995; 2013).

This study provides the first systematic description of maize-associated fungal communities across these endemic zones, offering a novel ecological basis for disease geography. Our results demonstrate that systematic differences in fungal community richness, taxonomic composition, and predicted biosynthetic capacity align closely with distinct disease settings. KD-endemic regions exhibited the highest fungal diversity and an enrichment of saprotrophic and symbiotrophic taxa typically associated with cool, humid, or transitional climates (Anthony et al. 2024; Lepinay et al. 2024). In contrast, KBD-endemic regions were dominated by fungi adapted to colder, persistently moist environments, including cellulose- and wood-degrading genera such as *Russula* and *Chaetomium*. Notably, KD–KBD co-endemic regions displayed the greatest functional breadth, characterized by the highest diversity and abundance of predicted biosynthetic gene clusters.

These fungal assemblages mirror global patterns of climate-driven structuring in cereal-associated mycobiomes (Tedersoo et al. 2014). Warm and humid agroecosystems typically favor *Aspergillus* and *Fusarium* species capable of producing aflatoxins and fumonisins, whereas temperate and cold climates favor *F. graminearum*, *F. culmorum*, *Claviceps purpurea*, and psychrotolerant fungi such as *Microdochium* and *Penicillium* spp.(Casu et al. 2024; Tsers et al. 2023). The fungal profiles observed in KD- and KBD-associated maize are consistent with this framework, suggesting that regional temperature and humidity regimes—rather than geographic location alone—are the primary determinants of fungal colonization and metabolic potential (Xu et al. 2025).

The ecological primacy of climate is further supported by the persistence of region-specific fungal signatures following controlled incubation under standardized conditions (Liu and Van der Fels-Klerx 2021). Despite uniform experimental settings, fungal communities retained clear distinctions corresponding to their regions of origin, highlighting the importance of resident fungal inocula and local adaptation (Khalaf et al. 2021; Wang et al. 2023). These findings indicate that climatic influences on fungal ecology extend beyond in situ colonization and continue to shape community dynamics during grain storage.

Taken together, these findings support a multifactorial etiological framework in which selenium deficiency acts as a host-level susceptibility factor, while climate-driven fungal community composition determines the nature and magnitude of dietary mycotoxin exposure. Selenium deficiency may impair antioxidant defenses and exacerbate oxidative stress, increasing vulnerability to fungal secondary metabolites (Bhattarai et al. 2024). Climatic conditions and post-harvest storage environments further shape fungal community dynamics, thereby influencing disease risk. This integrated model provides a coherent explanation for the longstanding epidemiological paradoxes associated with these “sister diseases.”

While this study establishes a comprehensive taxonomic and functional baseline, some limitations remain. Functional inference and BGC prediction were based on taxonomic annotation rather than direct metabolite measurement; future targeted mycotoxin analyses are needed to confirm exact exposure levels. Additionally, the cross-sectional design precludes an assessment of seasonal variation, and the role of the wider grain microbiome remains to be explored.

## 5. Conclusion

In conclusion, this study demonstrates that maize-associated fungal communities possess distinct taxonomic, functional, and metabolic signatures across KD- and KBD-endemic regions. By integrating ecological, climatic, and nutritional perspectives, our findings extend beyond the traditional low-selenium hypothesis and identify climate-sensitive fungal ecology as a critical environmental determinant of disease risk. These results provide a necessary foundation for future longitudinal studies and direct mycotoxin quantification to elucidate the causal pathways linking fungal metabolism to endemic disease prevention.

## Data Availability Statement

The raw sequence data generated in this study have been deposited in the NCBI Sequence Read Archive (SRA) database under accession number PRJNA1369992.

## Author Contributions

Yingxue Wang and Kunyu Zhang performed laboratory incubations and DNA extractions and drafted the original manuscript. Kunyu Zhang, Yu Sun, and Likun Yang conducted the literature review, performed bioinformatic analyses, and prepared the figures. Jinwu Yang and Xiaocheng Wang provided project administration and coordination. Yihong Wan, Guoping Xi, and Liuliu Guo were responsible for maize sample collection. Shuqiu Sun secured funding, conceived and designed the study, and critically revised the manuscript. All authors read and approved the final manuscript.

## Ethics Statement

Maize samples were collected from local households with the informed consent of the residents. The study did not involve human participants or vertebrate animals, and therefore no additional ethical approval was required.

## Supporting Information

Supplementary Table 1: Meteorological data including average temperatures and humidity for the study sites during the sample collection season.

Supplementary Figure 1: Fungal alpha and beta diversity across the seven individual sampling sites.

Supplementary Figure 2: Taxonomic composition of fungal communities at phylum and genus levels across the seven individual sampling sites.

Supplementary Figure 3: Relative abundances of mixed fungal trophic modes across the four disease regions.

## Reference

Allander, E., 1994. Kashin-beck disease. An analysis of research and public health activities based on a bibliography 1849-1992. Scand J Rheumatol Suppl 99, 1–36. 10.3109/03009749409117126.

Anthony, M.A., Tedersoo, L., De Vos, B., Croisé, L., Meesenburg, H., Wagner, M., Andreae, H., Jacob, F., Lech, P., Kowalska, A., Greve, M., Popova, G., Frey, B., Gessler, A., Schaub, M., Ferretti, M., Waldner, P., Calatayud, V., Canullo, R., Papitto, G., Marinšek, A., Ingerslev, M., Vesterdal, L., Rautio, P., Meissner, H., Timmermann, V., Dettwiler, M., Eickenscheidt, N., Schmitz, A., Van Tiel, N., Crowther, T.W., Averill, C., 2024. Fungal community composition predicts forest carbon storage at a continental scale. Nat Commun 15, 2385. 10.1038/s41467-024-46792-w.

Bhattarai, U., Xu, R., He, X., Pan, L., Niu, Z., Wang, D., Zeng, H., Chen, J.X., Clemmer, J.S., Chen, Y., 2024. High selenium diet attenuates pressure overload-induced cardiopulmonary oxidative stress, inflammation, and heart failure. Redox Biol 76, 103325. 10.1016/j.redox.2024.103325.

Casu, A., Camardo Leggieri, M., Toscano, P., Battilani, P., 2024. Changing climate, shifting mycotoxins: A comprehensive review of climate change impact on mycotoxin contamination. Compr Rev Food Sci Food Saf 23, e13323. 10.1111/1541-4337.13323.

Chasseur, C., Suetens, C., Nolard, N., Begaux, F., Haubruge, E., 1997. Fungal contamination in barley and kashin-beck disease in tibet. Lancet 350, 1074. 10.1016/s0140-6736(05)70453-0.

Chen, J., 2012. An original discovery: Selenium deficiency and keshan disease (an endemic heart disease). Asia Pac J Clin Nutr 21, 320–326.

Fang, H., Guo, X., Farooq, U., Xia, C., Dong, R., 2012. Development and validation of a quality of life instrument for kashin-beck disease: An endemic osteoarthritis in china. Osteoarthritis Cartilage 20, 630–637. 10.1016/j.joca.2012.03.004.

Guo, K., 1986. A review of research on the mycotoxin poisoning hypothesis for the etiology of keshan disease over the past thirty years. J Med Res 10, 289–294.

Khalaf, E.M., Shrestha, A., Rinne, J., Lynch, M.D.J., Shearer, C.R., Limay-Rios, V., Reid, L.M., Raizada, M.N., 2021. Transmitting silks of maize have a complex and dynamic microbiome. Sci Rep 11, 13215. 10.1038/s41598-021-92648-4.

Lei, C., Niu, X., Ma, X., Wei, J., 2011. Is selenium deficiency really the cause of keshan disease? Environ Geochem Health 33, 183–188. 10.1007/s10653-010-9331-9.

Lepinay, C., Větrovský, T., Chytrý, M., Dřevojan, P., Fajmon, K., Cajthaml, T., Kohout, P., Baldrian, P., 2024. Effect of plant communities on bacterial and fungal communities in a central european grassland. Environ Microbiome 19, 42. 10.1186/s40793-024-00583-4.

Li, G.S., Wang, F., Kang, D., Li, C., 1985. Keshan disease: An endemic cardiomyopathy in china. Hum Pathol 16, 602–609. 10.1016/s0046-8177(85)80110-6.

Li, H., Xing, Z., Dong, H., Qi, F., Yu, Q., Li, J., Jiang, H., Wang, C., Li, J., Zhang, B., Yu, J., 2026. T-2 toxin exacerbates chondrocyte extracellular matrix degradation potentially through yap/nlrp3/gsdmd-mediated pyroptosis. Int Immunopharmacol 168, 115890. 10.1016/j.intimp.2025.115890.

Lin, X., Liu, H., Qiao, L., Deng, H., Bao, M., Yang, Z., He, Y., Xiang, R., He, H., Han, J., 2024. Chondrocyte autophagy mediated by t-2 toxin via akt/tsc/rheb/mtor signaling pathway and protective effect of csa-senp. Osteoarthritis Cartilage 32, 1283–1294. 10.1016/j.joca.2024.05.007.

Liu, C., Van der Fels-Klerx, H.J., 2021. Quantitative modeling of climate change impacts on mycotoxins in cereals: A review. Toxins (Basel) 13. 10.3390/toxins13040276.

Lu, Q., Hu, S., Guo, P., Zhu, X., Ren, Z., Wu, Q., Wang, X., 2021. Ppar-γ with its anti-fibrotic action could serve as an effective therapeutic target in t-2 toxin-induced cardiac fibrosis of rats. Food Chem Toxicol 152, 112183. 10.1016/j.fct.2021.112183.

Prabhu, K.S., Lei, X.G., 2016. Selenium. Adv Nutr 7, 415–417. 10.3945/an.115.010785.

Sun, L., Cui, S., Deng, Q., Liu, H., Cao, Y., Wang, S., Yu, J., 2019. Selenium content and/or t-2 toxin contamination of cereals, soil, and children’s hair in some areas of heilongjiang and gansu provinces, china. Biol Trace Elem Res 191, 294–299. 10.1007/s12011-018-1620-7.

Sun, S.Q., 2010. Chronic exposure to cereal mycotoxin likely citreoviridin may be a trigger for keshan disease mainly through oxidative stress mechanism. Med Hypotheses 74, 841–842. 10.1016/j.mehy.2009.11.043.

Sun, S.Q., 2012. Etiological research on keshan disease: Transition from hypothesis debate to clue investigation. Chinese Journal of Endemiology 31, 241–244. 10.3760/cma.j.issn.1000-4955.2012.03.002.

Sun, S.Q., Ji, T., & Zhang, J. N, 2018. A brief history and enlightenment of the etiological research on keshan disease. Chinese Journal of Endemiology 37, 345–350. 10.3760/cma.j.issn.2095-4255.2018.05.001.

Tedersoo, L., Bahram, M., Põlme, S., Kõljalg, U., Yorou, N.S., Wijesundera, R., Villarreal Ruiz, L., Vasco-Palacios, A.M., Thu, P.Q., Suija, A., Smith, M.E., Sharp, C., Saluveer, E., Saitta, A., Rosas, M., Riit, T., Ratkowsky, D., Pritsch, K., Põldmaa, K., Piepenbring, M., Phosri, C., Peterson, M., Parts, K., Pärtel, K., Otsing, E., Nouhra, E., Njouonkou, A.L., Nilsson, R.H., Morgado, L.N., Mayor, J., May, T.W., Majuakim, L., Lodge, D.J., Lee, S.S., Larsson, K.H., Kohout, P., Hosaka, K., Hiiesalu, I., Henkel, T.W., Harend, H., Guo, L.D., Greslebin, A., Grelet, G., Geml, J., Gates, G., Dunstan, W., Dunk, C., Drenkhan, R., Dearnaley, J., De Kesel, A., Dang, T., Chen, X., Buegger, F., Brearley, F.Q., Bonito, G., Anslan, S., Abell, S., Abarenkov, K., 2014. Fungal biogeography. Global diversity and geography of soil fungi. Science 346, 1256688. 10.1126/science.1256688.

Tsers, I., Marenina, E., Meshcherov, A., Petrova, O., Gogoleva, O., Tkachenko, A., Gogoleva, N., Gogolev, Y., Potapenko, E., Muraeva, O., Ponomareva, M., Korzun, V., Gorshkov, V., 2023. First genome-scale insights into the virulence of the snow mold causal fungus microdochium nivale. IMA Fungus 14, 2. 10.1186/s43008-022-00107-0.

Wang, K., Yu, J., Liu, H., Liu, Y., Liu, N., Cao, Y., Zhang, X., Sun, D., 2020. Endemic kashin-beck disease: A food-sourced osteoarthropathy. Semin Arthritis Rheum 50, 366–372. 10.1016/j.semarthrit.2019.07.014.

Wang, M., Wang, C., Yu, Z., Wang, H., Wu, C., Masoudi, A., Liu, J., 2023. Fungal diversities and community assembly processes show different biogeographical patterns in forest and grassland soil ecosystems. Front Microbiol 14, 1036905. 10.3389/fmicb.2023.1036905.

Wang, X., Wang, S., He, S., Zhang, F., Tan, W., Lei, Y., Yu, H., Li, Z., Ning, Y., Xiang, Y., Guo, X., 2013. Comparing gene expression profiles of kashin-beck and keshan diseases occurring within the same endemic areas of china. Sci China Life Sci 56, 797–803. 10.1007/s11427-013-4495-z.

Xu, J., Zhang, Y., Ren, J., Kong, Q., 2025. An atoxigenic aspergillus flavus pa67 from shandong province exhibits potential in biocontrol against toxigenic aspergillus flavus, sclerotium rolfsii, and fusarium proliferatum. Int J Food Microbiol 426, 110918. 10.1016/j.ijfoodmicro.2024.110918.

Yan, H., Sun, J., Fu, X., Ye, J., Wang, W., Cao, J., Ji, J., Sun, X., 2026. Climate change: An inevitable factor in reshaping the contamination level of fungi and mycotoxins. Compr Rev Food Sci Food Saf 25, e70354. 10.1111/1541-4337.70354.

Yang, J.B., 1995. Research report on the etiology of kashin-beck disease. Chinese Journal of Endemiology. 10.3969/j.issn.1001-1889.2005.06.010.

Yang, J.B., 2002. Mycotoxins and human diseases. Chinese Journal of Endemiology 21, 314–317. 10.3760/cma.j.issn.1000-4955.2002.04.027.

Yang, J.B., 2013. Geographical distribution characteristics and causes of keshan disease and kashin-beck disease. Chin J Endemiol 32, 4–6. 10.3760/cma.j.issn.2095-4255.2013.01.002.

Yang, J.B., & Yang, Q. H., 2000. Research on the etiology of keshan disease. Chinese Journal of Endemiology 19, 350–355. CNKI:SUN:ZDFB.0.2000-05-012.

Yu, F.F., Zuo, J., Sun, L., Yu, S.Y., Lei, X.L., Zhu, J.H., Zhou, G.Y., Guo, X., Ba, Y., 2022. Animal models of kashin-beck disease exposed to environmental risk factors: Methods and comparisons. Ecotoxicol Environ Saf 234, 113419. 10.1016/j.ecoenv.2022.113419.

Zhang, D., Wei, C., Yu, J. N., 2012. Effect of citreoviridin on myocardial morphology and structure in rats fed with low selenium and low protein diet. Chinese Journal of Endemiology 31, 382–385. 10.3760/cma.j.issn.1000-4955.2012.04.011.

Zou, K., Liu, G., Wu, T., Du, L., 2009. Selenium for preventing kashin-beck osteoarthropathy in children: A meta-analysis. Osteoarthritis Cartilage 17, 144–151. 10.1016/j.joca.2008.06.011.

